# Unveiling the antiviral capabilities of targeting Human Dihydroorotate Dehydrogenase against SARS-CoV-2

**DOI:** 10.1101/2023.10.04.560875

**Authors:** Aline D. Purificação, Sabrina S. Mendonça, Luiza V. Cruz, Carolina Q. Sacramento, Jairo R. Temerozo, Natalia Fintelman-Rodrigues, Caroline Souza de Freitas, Bruna F. Godoi, Miguel M. Vaidergorn, Juliana Almeida Leite, Luis Carlos Salazar Alvarez, Murillo V. Freitas, Meryck F. B. Silva, Bianca A. Martin, Renata F. V. Lopez, Bruno J. Neves, Fabio T. M. Costa, Thiago M. L. Souza, Flavio da S. Emery, Carolina Horta Andrade, M. Cristina Nonato

**Affiliations:** Protein Crystallography Laboratory, Department of Biomolecular Sciences, School of Pharmaceutical Sciences at Ribeirao Preto, University of São Paulo, Ribeirão Preto, SP, Brazil; Center for the Research and Advancement in Fragments and molecular Targets (CRAFT), School of Pharmaceutical Sciences at Ribeirao Preto, University of São Paulo, Ribeirão Preto, SP, Brazil; Laboratory for Molecular Modeling and Drug Design (LabMol), Faculty of Pharmacy, Universidade Federal de Goiás, Goiânia, 74605-170, GO, Brazil; Laboratory of Immunopharmacology, Oswaldo Cruz Institute, Fiocruz, Rio de Janeiro, RJ, Brazil; National Institute for Science and Technology on Innovation in Diseases of Neglected Populations (INCT/IDPN), Center for Technological Development in Health (CDTS), Fiocruz, Rio de Janeiro, RJ, Brazil; National Institute for Science and Technology on Neuroimmunomodulation, Oswaldo Cruz Institute, Fiocruz, Rio de Janeiro, RJ, Brazil; Laboratory of Heterocyclic and Medicinal Chemistry (QHeteM), Department of Pharmaceutical Sciences, School of Pharmaceutical Sciences at Ribeirao Preto, Ribeirao Preto, SP, Brazil; Laboratory of Tropical Diseases, Dep. Genetics, Evolution, Microbiology and Immunology, Institute of Biology, Unicamp, Campinas, SP, Brazil; Innovation Center in nanostructured systems and topical administration (NanoTop), School of Pharmaceutical Sciences at Ribeirao Preto, University of São Paulo, Ribeirão Preto, SP, Brazil; Center for Excellence in Artificial Intelligence (CEIA), Institute of Informatics, Universidade Federal de Goiás, Goiânia, 74605-170, GO, Brazil

## Abstract

The urgent need for effective treatments against emerging viral diseases, driven by drug-resistant strains and new viral variants, remains critical. We focus on inhibiting the human dihydroorotate dehydrogenase (*Hs*DHODH), one of the enzymes in charge of pyrimidine nucleotide synthesis. This strategy could impede viral replication without provoking resistance. We evaluated quinone-based compounds, discovering potent *Hs*DHODH inhibition (low nanomolar IC_50_) and promising in vitro anti-SARS-CoV-2 activity (low micromolar EC_50_). These compounds exhibited low toxicity, indicating potential for further development. Additionally, we employed computational tools like molecular docking and QSAR models to analyze protein-ligand interactions. These findings represent a significant step forward in the search for effective antiviral treatments and have great potential to impact the development of new broad-spectrum antiviral drugs.

## INTRODUCTION

Since the first evidence of COVID-19 in December 2019, the world has suffered the devastating consequences of the pandemic caused by SARS-CoV-2, resulting in more than 750 million confirmed cases and almost 7 million deaths as of March 29, 2023 (1).

In this scenario, a worldwide race to find therapeutic options for COVID-19 started, using both drug discovery and drug repositioning. Drug repositioning allowed the approval of different small-molecule drugs, generally broad-spectrum antivirals such as Remdesivir (2–4) that was initially developed to treat the Ebola Virus. Another commercially available drug for rheumatoid arthritis, Leflunomide, has been extensively studied for its potential anti-SARS-CoV-2 action (5–8). Leflunomide is a prodrug, and its active metabolite, teriflunomide, is a potent inhibitor of the human enzyme dihydroorotate dehydrogenase (DHODH), which participates in the *de novo* synthesis of pyrimidine nucleotides.

The *de novo* synthesis of pyrimidine nucleotides is a pathway conserved across bacteria (9,10), protozoan (11), animal (12–14), and plants (15), supplying the basic building blocks for the nucleic acids DNA and RNA. Recently, this pathway has been studied as a target for broad-spectrum antivirals since viruses rely on specific host factors to complete its lifecycle (16).

Cells with high replication rates, such as those involved in viral replication, have a greater demand for nucleotides compared to slowly replicating cells, including most adult human cells (17). Thus, targeting key enzymes involved in host nucleotide biosynthesis represents a promising broad-spectrum antiviral strategy (18,19). By inhibiting these enzymes, we can disrupt the availability of nucleotides crucial for viral replication, thereby impeding the viral lifecycle and proliferation within the host cell.

DHODH is an enzyme involved in the *de novo* biosynthesis of pyrimidines, responsible for the fourth step, the only redox reaction, and a rate-limiting step in the pathway (20). Located on the outer surface of the inner membrane of mitochondria, human DHODH (*Hs*DHODH) converts dihydroorotate to orotate, subsequently utilized to produce uridine monophosphate (UMP) and, finally, pyrimidine nucleotide (20–22) (FIGURE 1).

**FIGURE 1:**
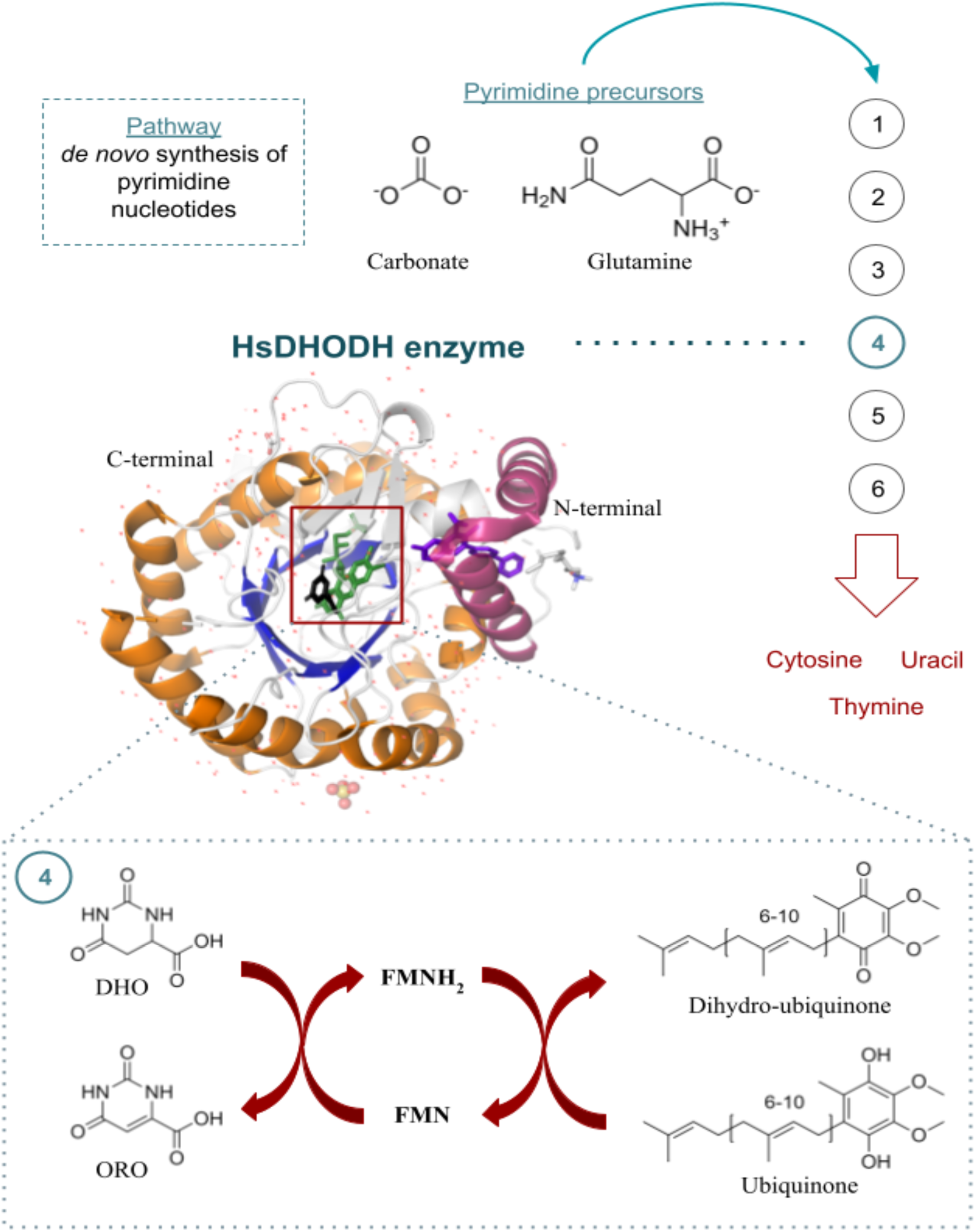
*Hs*DHODH is part of the *de novo* pyrimidine biosynthesis. The synthesis of pyrimidine nucleotides relies on the precursors carbonate and glutamine and involves six steps catalyzed by carbamoyl phosphate synthetase (step 1), aspartate transcarbamylase (step 2), dihydroorotase (step 3), dihydroorotate dehydrogenase (step 4), and the bifunctional enzyme UMP synthase, which is involved in the final two steps (22). The *Hs*DHODH enzyme plays a key role in the fourth step of the synthesis, catalyzing an oxi-reduction reaction via a ping-pong mechanism that converts dihydroorotate (DHO) to orotate (ORO) using the monoflavin FMN as a cofactor, which is then recovered by the ubiquinone molecule. This pathway is essential for maintaining an adequate supply of intracellular pyrimidine nucleotides.

The inhibition of pyrimidine biosynthesis has been utilized as a therapeutic target in various fields, including rheumatology (23), oncology (24,25), immunological disorders (26), and infectious diseases (5,16,27–35), due to its immunomodulatory properties. In this study, we evaluated the inhibitory potential of a small group of quinone-based compounds against *Hs*DHODH and their in vitro anti-SARS-CoV-2 activity. Furthermore, we employed computational approaches, such as molecular docking and quantitative structure-activity relationship (QSAR) models, to analyze the protein-ligand interactions and identify the fragments responsible for the antiviral activity. These findings provide valuable insights for future hit-to-lead optimization campaigns.

## RESULTS AND DISCUSSION

In our previous work, we designed several quinoidal analogs of atovaquone and evaluated their effectiveness against *Schistosoma mansoni* DHODH (*Sm*DHODH), using *Hs*DHODH as a reference to assess selectivity for the parasite enzyme (36). While some compounds displayed substantial activity against *Sm*DHODH, they lacked selectivity and also inhibited *Hs*DHODH with high potency (36). Consequently, in this study, we revalidated seven of these potent naphthoquinoidal *Hs*DHODH inhibitors in terms of their enzymatic inhibition and evaluated their experimental anti-SARS-CoV-2 and cytotoxicity data for the first time (TABLE 1, FIGURE 2). As experimental controls, we used the *Hs*DHODH inhibitors brequinar, teriflunomide, and ML390, which all fell within the nanomolar range (FIGURE 2). The tested compounds belong to the 2-hydroxynaphthoquinone (NQ) family and can be classified into two groups based on their chemical structure. The first group comprises a 3-alkyl-substituted NQ, which includes lapachol (QHM001) with an isoprenyl chain and reduced lapachol (QHM020) with a 3-isopentyl 3-substituted-NQ. The second group is a series of 3-anilines-NQ (FIGURE 2).

**TABLE 1.** The activity of the tested quinones regarding enzyme inhibition, anti-SARS-CoV-2 activity, and cytotoxicity. Table 1 presents the in vitro values for enzyme inhibition (IC_50_ in *Hs*DHODH), anti-SARS-CoV-2 activity (EC_50_ in infected Calu-3 cells), and cytotoxicity (CC_50_ in both HepG2 and Calu-3 cells) of the tested quinones and experimental controls. The selective index for each compound is also reported, representing the correlation between its anti-SARS-CoV-2 activity and HepG2 cytotoxicity.

| | $IC_{50}$ nM | $EC_{50}$ μM | $CC_{50}$ μM<br>(HepG2) | $CC_{50}$ μM<br>(Calu-3) | Selective<br>index |
| --- | --- | --- | --- | --- | --- |
| QHM001 | $48 \pm 4$ | $1.5 \pm 0.1$ | $71 \pm 9$ | $67 \pm 2$ | $47 \pm 7$ |
| QHM011 | $685 \pm 53$ | $2.3 \pm 0.3$ | $176 \pm 23$ | $77 \pm 2$ | $77 \pm 14$ |
| QHM020 | $293 \pm 32$ | $1.8 \pm 0.2$ | $65 \pm 12$ | $75 \pm 2$ | $35 \pm 7$ |
| QHM034* | $365 \pm 23$ | ND | $324 \pm 86$ | ND | ND |
| QHM110 | $101 \pm 3$ | $1.5 \pm 0.2$ | $178 \pm 35$ | $48 \pm 2$ | $117 \pm 27$ |
| QHM230 | $468 \pm 33$ | $1.2 \pm 0.2$ | $141 \pm 33$ | $67 \pm 2$ | $114 \pm 32$ |
| QHM297** | $307 \pm 54$ | ND | $17 \pm 4$ | ND | ND |
| ML390 | $198 \pm 14$ | ND | $16 \pm 5$ | $91 \pm 2$ | ND |
| Brequinar | $5.5 \pm 0.5$ | $0.8 \pm 0.1$ | $17 \pm 4$ | $48 \pm 2$ | $20 \pm 5$ |
| Teriflunomide | $145 \pm 4$ | ND | $164 \pm 18$ | $59 \pm 2$ | ND |
| Remdesivir | NA | $0.08 \pm 0.02$ | NA | NA | NA |

All values are displayed as mean ± standard error; ND not disponible data; NA not applicable data. \* chemically unstable. \*\* due to the high toxicity observed in HepG2 cell lines, this compound was not assessed in Calu-3 cell experiments.

**FIGURE 2:**
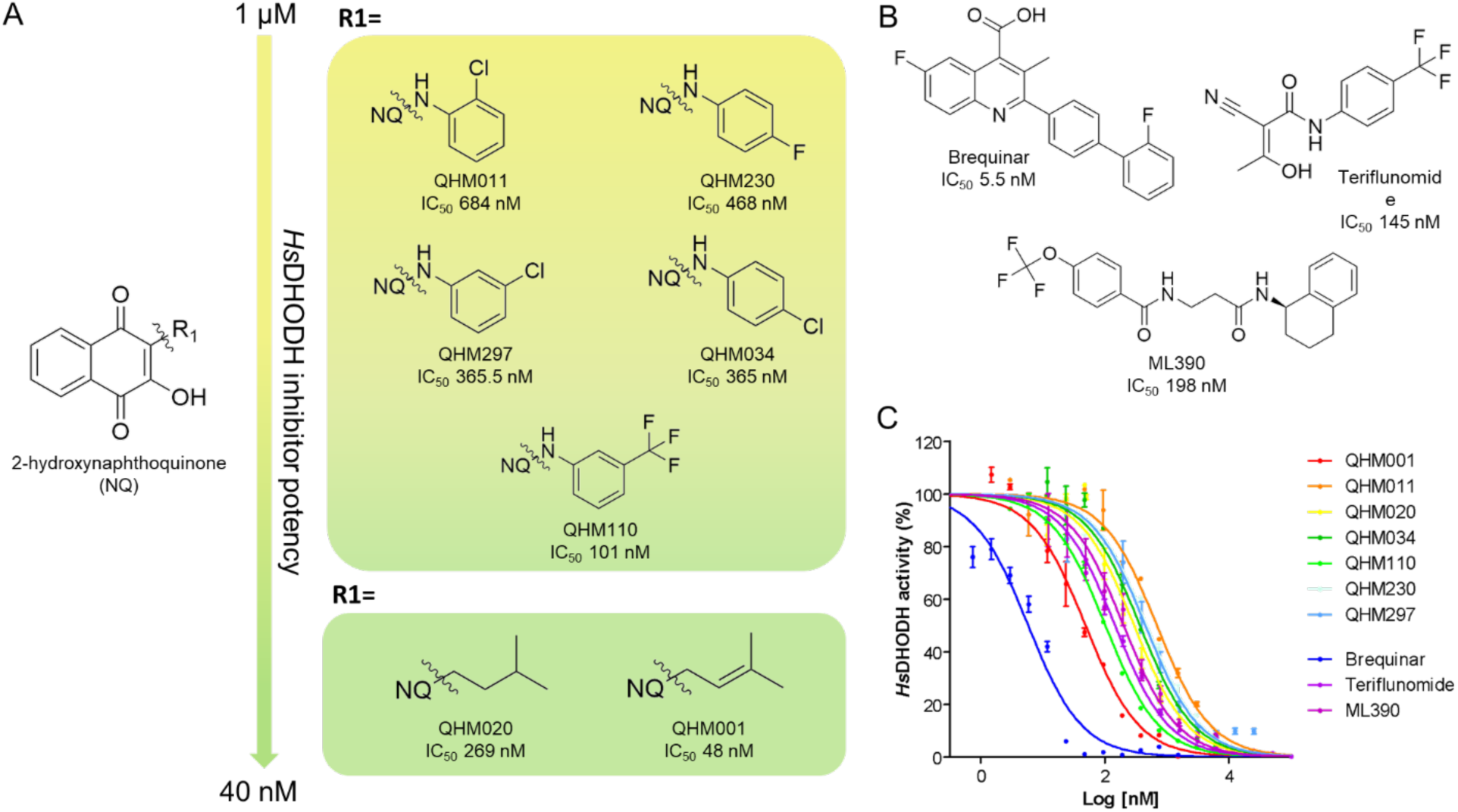
Napthoquinones are potent *Hs*DHODH inhibitors. The structures and dose-response curves of enzymatic inhibition for the tested compounds are shown. A) The structures of the 3-alkyl-substituted-NQ and 3-isopentyl 3-substitute-NQ series are sorted by potency against *Hs*DHODH, ranging from 1µM to 40 nM. B) The chemical structures of the commercial and clinical *Hs*DHODH inhibitors, including Brequinar, Teriflunomide, and ML390, are shown. C) The IC_50_ curves for all the quinones tested in this study, as well as the experimental controls, are presented.

The most potent compound tested was lapachol (QHM001), which had an IC_50_ value of 48 nM. Interestingly, when the isoprenyl chain of lapachol was reduced (QHM020), the inhibitory potential decreased by approximately five-fold (TABLE 1, FIGURE 2) (37). The isopentyl portion of QHM020 is more flexible, less lipophilic, and less electronegative than the allyl group. The difference in IC_50_ values for QHM001 and QHM020 suggests that these properties may guide the potency of the compounds in *Hs*DHODH. However, the similarity in IC_50_ values for QHM020 and compounds of the 3-anilines-NQ series, such as QHM110, QHM034, and QHM297, which have a bulkier R1 (FIGURE 2), strongly suggests that differences in molecular volume do not impact enzyme inhibition.

Upon examination of compounds QHM034, QHM297, and QHM230, it becomes evident that chloride and fluoride in para- or meta-substituted anilines have similar effects on enzyme inhibition (TABLE 1, FIGURE 2) (37). Thus, differences in the physicochemical properties of halogens, such as electronegativity and atomic radius, do not appear to impact enzyme inhibition. However, the halogen in the ortho position is less favorable (QHM011) compared to the meta- and para-substituted benzene (QHM230, QHM297, QHM034).

Interestingly, replacing a halogen with a bulky trifluoromethyl group (QHM110) resulted in a significant increase in potency (TABLE 1, FIGURE 2) (37). Notably, QHM110 shares similarities with teriflunomide (IC_50_ 145 nM), which also possesses the trifluoromethyl group in the para-benzene position. This similarity is evident in the superposition of both structures in *Hs*DHODH, as illustrated in FIGURE S1.

To better understand how these compounds interact with *Hs*DHODH and elicit structure-activity relationships (SAR), we conducted molecular docking for each compound with this enzyme. Additionally, we calculated the electrostatic potential map for the quinones (FIGURE 3). The Glide XP Scores, Ligand Efficiency (LE), and MMGBSA scores can be found in TABLE S1, while the 3D diagrams showing the superposition between the docked compounds in *Hs*DHODH and the crystal structure of *Hs*DHODH co-crystallized with brequinar (PDB ID: 1D3G) and teriflunomide (PDB ID: 1D3H) are available in FIGURE S2.

**FIGURE 3:**
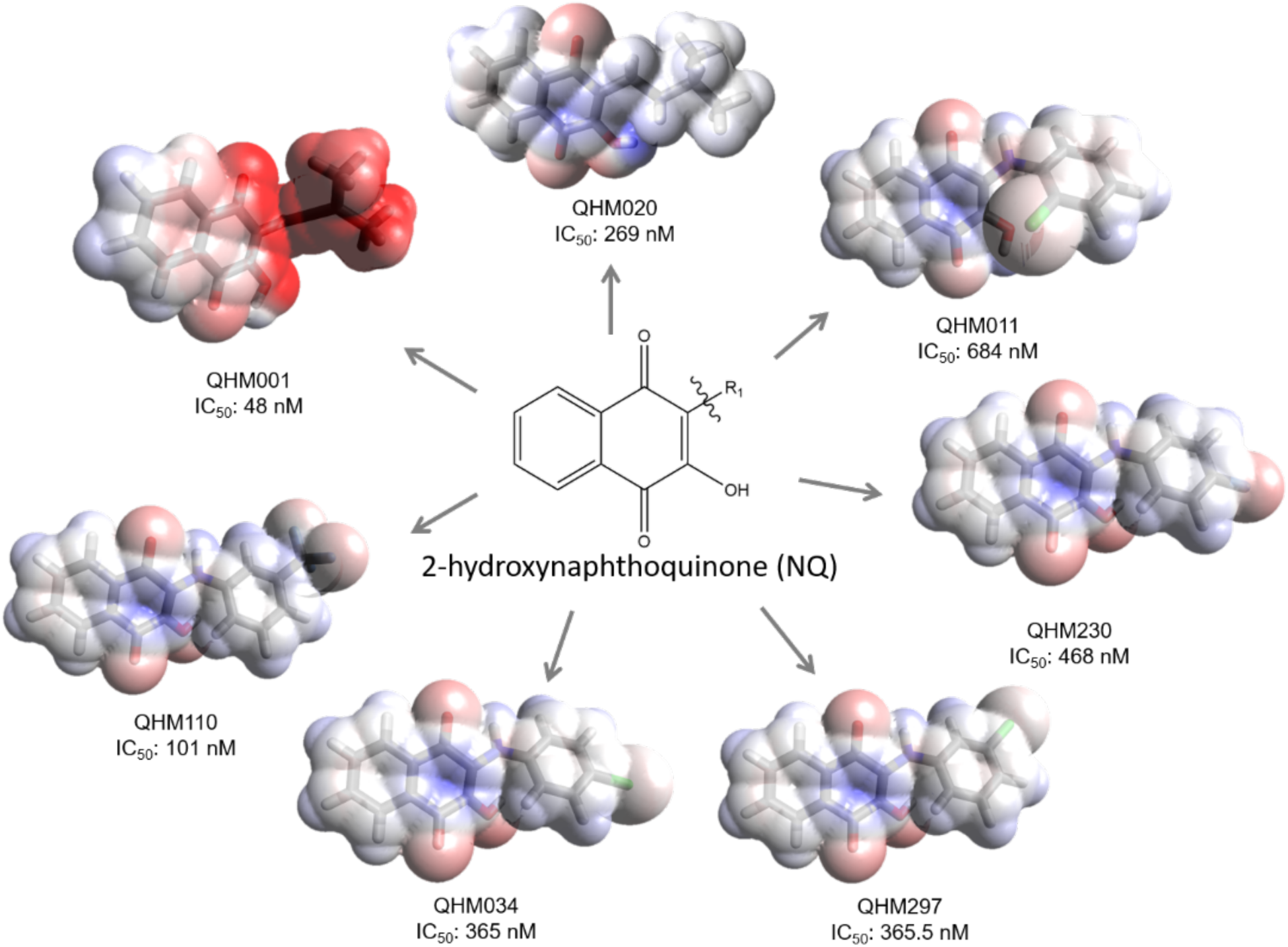
Electrostatic potential map of tested quinones. The diagram illustrates the electrostatic differences between compounds and their relationship with IC_50_ values. Compounds are presented in sticks models, with carbon atoms in gray, hydrogen in white, oxygen in red, nitrogen in blue, chlorine in green and fluorine in light blue. The electrostatic potential is represented as the van der Walls radius of the atoms, where red, blue, and white indicate negative, positive, and neutral potential, respectively.

In addition to being more potent, QHM001 also exhibits higher electronic density potential, primarily concentrated in the 2-hydroxy-p-quinone portion, where the double bond is located (FIGURE 3). This is not observed in QHM020, which lacks the double bond and therefore reduces the electronic density potential in this region. Notably, the most potent compound of the 3-anilines-NQ series, QHM110, also possesses a portion with high electronic density potential in the CF_3_ position. To emphasize the significance of higher electron density groups in the isoprenyl or CF_3_ position, which may have contributed to stronger interactions with the enzyme in the hydrophobic region, we have included 3D representations of docked QHM001 and QHM110 with the molecular electrostatic surface of the protein in FIGURE S2. Interestingly, the higher electron density of the hydroxyquinone moiety in QHM001, compared to the other compounds tested, could be linked to stronger polar interactions with the molecular target, potentially resulting in higher inhibitory potency.

Molecular docking calculations indicate a good fit for all quinone compounds. Their superposition with brequinar and teriflunomide reveals a very similar interaction model for the docked compounds, as seen in brequinar (FIGURE S1). According to Baumgartner’s (38) nomination, all the quinones interact with subsites 1 (M43, L46, A55), 2 (R136), 3 (Y356), and 4 (V134, V143), and the same is observed for brequinar (PBDID 1D3G) (39). However, brequinar has additional residues involved in the interaction in the subsites 1 (Y38, L42, L68, F58, F98, and M111) and subsite 5 (L67), a fact that can be related to the higher potency of brequinar (IC_50_ 5.5 nM) compared to the tested quinones.

The quinones form an additional H-bond with T360 of *Hs*DHODH, mediated by a water molecule, which is not observed in brequinar and teriflunomide. This information suggests a chemical basis for the modulation of this target.

Since the quinones have a “brequinar-like” interaction pattern with the enzyme and considering the adverse effects observed in clinical trials with brequinar, we decided to assess the potential toxicity of these new molecules via *in vitro* cellular toxicity assays, which is a key point in drug discovery. Cytotoxicity assays were performed on two different cell lines. Naturally, we used Calu-3 cells, a pneumocyte lineage, to compare the antiviral effect, which was studied in this same cell type. Pneumocytes are the main cells affected during severe COVID-19. In addition, we also evaluated the cytotoxicity in cells of hepatic origin, the HepG-2 cell line. Considering our future interest in developing these analogs for potential oral use, these substances would reach the liver via the hepatic portal circulation. Therefore, cytotoxicity in liver cell lines was also evaluated. To that end, we used approved drug teriflunomide as experimental control. Brequinar exhibited the lowest CC_50_ value (17 µM), whereas teriflunomide showed higher CC_50_ value (164 µM). The quinones QHM297 (17 µM) and QHM001 (71 µM) also demonstrated high cytotoxicity. The other evaluated quinone-based compounds exhibited lower cytotoxicity than the approved drugs (TABLE 1). We then conducted anti-SARS-CoV-2 assays in Calu-3 cells infected with SARS-CoV-2 to determine the EC_50_ values for the *Hs*DHODH inhibitors. The most potent anti-SARS-CoV-2 quinone-based compound was QHM230, which was also the most toxic. All the tested compounds exhibited an EC_50_ in the low micromolar range (FIGURE S3). The calculated selective index indicated that all tested quinone compounds were more selective than brequinar, suggesting that these compounds may cause fewer side effects than those observed with brequinar in several clinical studies.

Although all docked quinones show a binding mode similar to brequinar, there are some differences in the interactions of each docked molecule. By analyzing the most potent quinone against *Hs*DHODH, QHM001, and the most selective anti-SARS-CoV-2 compound, QHM110, we observed that the main interactions are preserved (FIGURE 4). However, slight differences in the inhibitory site could be responsible for the contrast in biological assays. QHM001 (CC_50_ 71 µM) is much more cytotoxic than QHM110 (CC_50_ 178 µM), resulting in a selectivity difference of more than two times for the antiviral assays. Similar to the interaction pattern of brequinar compared to the tested quinones, QHM001 has more hydrophobic interactions (14 interactions: A59, M43, L46, F62, L58, A55, G47, H56, P52, T360, V134, V143, Y147, Y356) than QHM110 (12 interactions: A59, M43, L46, F62, A55, H46, P52, T360, V134, V143, Y356, L359), supporting the hypothesis that hydrophobic interactions are crucial for target modulation. However, the participation of hydrophobic interactions seems to be related to cytotoxicity, as brequinar makes more hydrophobic interactions than QHM001, which makes more hydrophobic interactions than QHM110. This pattern is also observed in cytotoxicity. This information could be valuable in guiding the synthesis of new molecules that are more promising in biological assays and potentially safer and more effective in the future.

**FIGURE 4:**
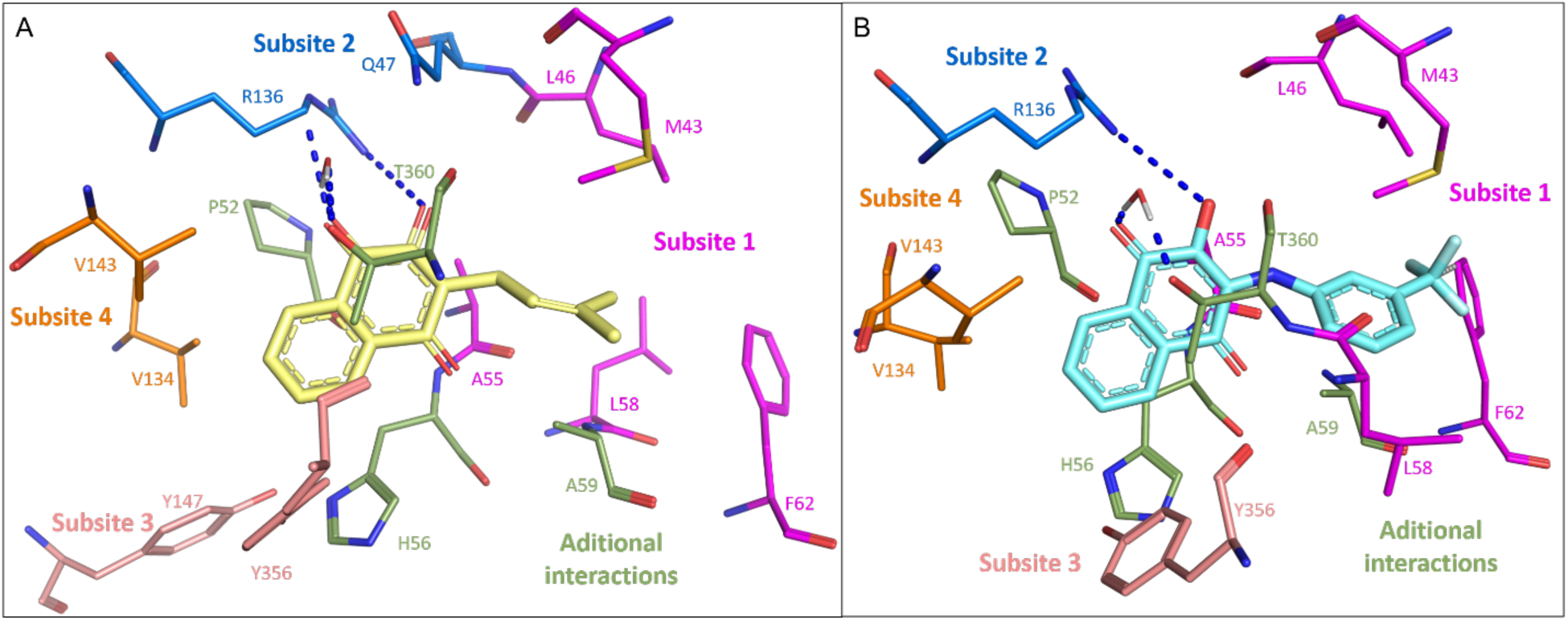
Quinones have the “brequinar-like” binding mode on *Hs*DHODH. Docked poses obtained for (A) QHM001 and (B) QHM110 (carbon atoms cyan). The subsites of the protein are colored according to Baumgartner’s (38) nomination, subsite 1 (magenta), subsite 2 (dark blue), subsite 3 (salmon), and subsite 4 (orange). The quinones have additional interactions with hydrophobic residues, colored in green.

To better understand the properties of our new brequinar-like molecules and prepare for upcoming pharmacokinetic studies, we analyzed their water solubility. Water solubility is an essential aspect of drug discovery, as it affects the dissolution rate and can decrease the absorption and bioavailability of the drug (40,41). According to the FDA’s biopharmaceutical classification system guide (42), a drug is highly soluble if its therapeutic dose is soluble in 250 mL, approximately one water glass, of an aqueous medium. Despite their similar molecular structures, the quinones exhibited different water solubility coefficients (TABLE S2). Interestingly, the quinones with the highest and lowest solubility coefficients were, respectively, QHM034 (100 μM) and QHM011 (3 μM), both of which are isomers of the chlorine-substituted aniline. Considering the administration of 1 mg of the synthesized compounds, their solubility should be higher than 4 μg/mL to be considered a highly soluble drug. Using this FDA criterion, only QHM011, QHM020 and QHM230 have low solubility (TABLE S2). All other quinones showed water solubility that could be modulated with simple strategies, such as preparing amorphous solid dispersions (43), making them promising candidates for drug discovery campaigns.

In this study, we developed QSAR models using machine learning and deep learning algorithms to classify untested compounds as active or inactive based on a public dataset of compounds evaluated for the cytopathic effect induced by SARS-CoV-2 infection in Vero E6 cells. These models provide a visual interpretation of the fragments responsible for antiviral activity.

Supplementary TABLE S3 provides complete statistical characteristics of the training, validation, and test sets for the generated models. FIGURE 5A shows the statistical characteristics of the best classification models on the test sets. The closer the accuracy (ACC), sensitivity (SE), and specificity (SP) values are to 1, the better the model performed. As shown in FIGURE 5A, the feedforward neural network (FFNN) architecture outperformed the models obtained by graph representations (i.e., message-passing neural network) and machine learning (i.e., Random Forest). Our FFNN model exhibited balanced SE (1.0 ± 0.0) and SP (0.94 ± 0.03) values, indicating that the model can accurately detect both active and inactive compounds. The FFNN model was developed using a simple architecture (FIGURE 5B) consisting of two fully connected hidden layers with 10 and 3 units. Both layers were configured with a non-linear Leaky ReLU activation function and dropout of 0.1 to prevent model overfitting.

**FIGURE 5:**
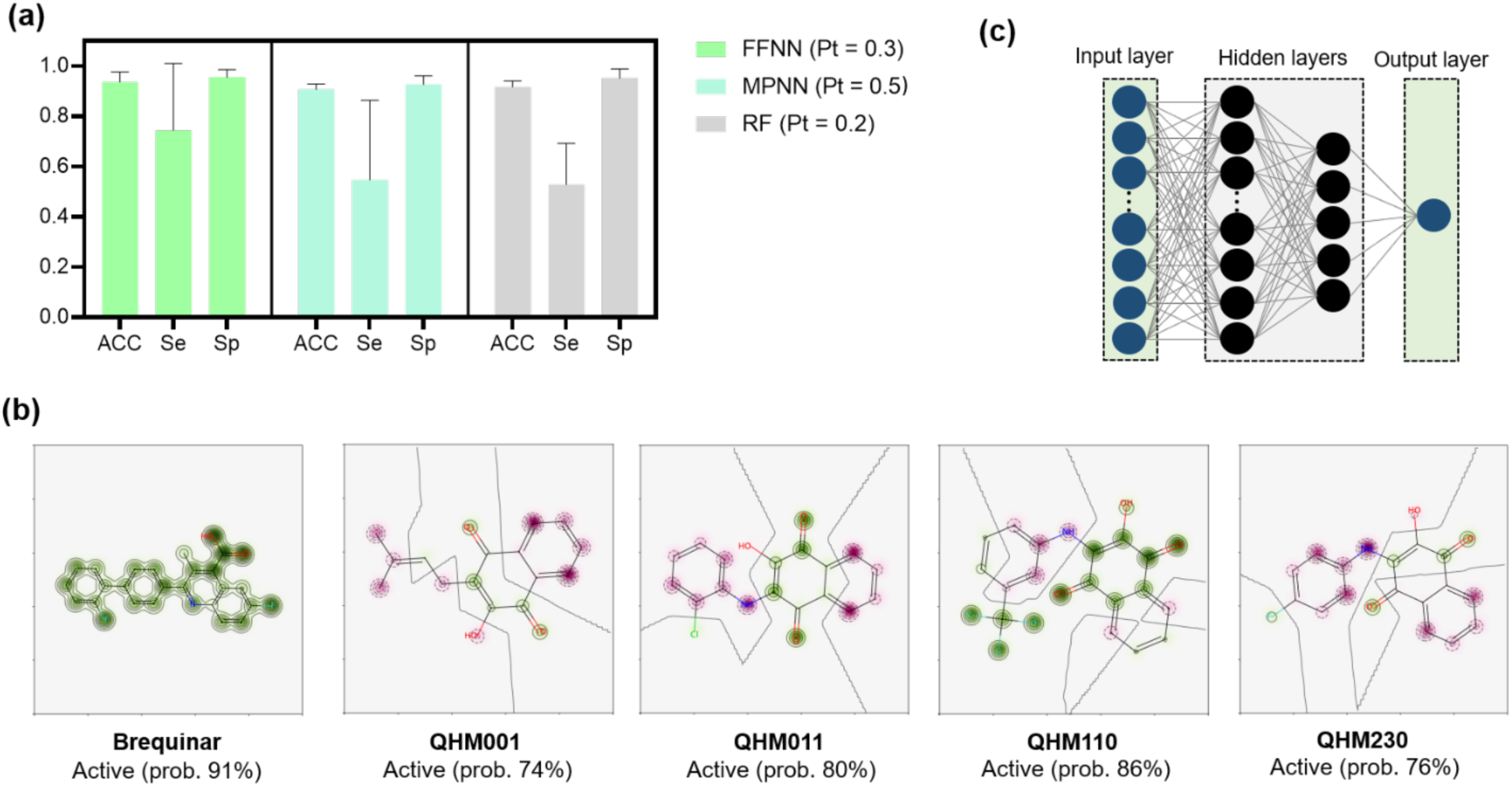
QSAR models for predicting compounds with antiviral activity. Panel (A) displays the statistical characteristics of the best classification models on the test sets, where higher values of ACC, SE, and SP indicate better performance. The optimal probability threshold (Pt) is also indicated. Panel (B) shows the FFNN architecture used for classifying compounds as active or inactive against SARS-CoV-2. Panel (C) depicts the predicted influence probability maps of structural fragments for the experimental hits and brequinar on the antiviral activity generated from the best FFNN model. Fragments that promote positive contributions to the antiviral activity are highlighted in green, while those that decrease it are highlighted in pink.

The compounds identified as the best experimental hits in the cell-based assays with SARS-CoV-2, including QHM001, QHM011, QHM110, and QHM230, as well as the positive control brequinar, were evaluated using the best FFNN classification model. The high predictive power of the model was confirmed as all the compounds in the study were correctly predicted as actives on SARS-CoV-2, with probabilities greater than 70%. Significant atomic and fragment features that prevent the cytopathic effect induced by SARS-CoV-2 were identified, providing structural insights for prospective hit-to-lead optimization. The results indicated that carbonyl and hydroxy groups of the 2-hydroxy-1,4-naphthoquinone moiety positively contribute to the activity, which supports their importance as hydrogen bond acceptors, as previously suggested by molecular docking analyses. These features are highlighted in green in FIGURE 5C, while fragments that decrease the activity are shown in pink.

Therefore, the FFNN classification models developed in this study could be potentially applied in future studies aimed at discovering new compounds with anti-SARS-CoV-2 activity. The highlighted structural fragments in green could also serve as valuable guidance for optimization in subsequent studies. Additionally, all models, data, and codes used in this research are available on GitHub (https://github.com/LabMolUFG/COVID).

## FINAL CONSIDERATIONS

The search for effective treatments against emerging and reemerging viral diseases has become increasingly urgent in recent years, as drug-resistant strains and new viral variants continue to emerge. In this context, our study makes a substantial contribution to the field of antiviral drug discovery, focusing on the activity against SARS-CoV-2, the virus responsible for the COVID-19 pandemic. By targeting the human dihydroorotate dehydrogenase enzyme, which plays a critical role in the *de novo* synthesis of pyrimidine nucleotides, we were able to inhibit viral replication and identify a series of naphthoquinoidal atovaquone analogs with potent anti-SARS-CoV-2 activity with low toxicity, suggesting their potential for further optimization and development. Additionally, we employed computational techniques like molecular docking and QSAR models using machine learning and deep learning algorithms to investigate the protein-ligand interactions and identify the specific fragments responsible for the observed antiviral activity.

Notably, our study demonstrates that targeting *Hs*DHODH represents a promising strategy for antiviral drug discovery, as this host enzyme is less likely to cause resistance and is not associated with possible virus mutations. Our study represents a significant step forward in the search for effective treatments against viral diseases, and our findings could significantly impact the development of new antiviral drugs.

## METHODOLOGY

### Inhibition assays

The inhibitory assays were performed for indirect measure of dichlorophenol-indophenol (DCIP) reduction as previously described (36) in a 96-well microplate reader, in a reaction buffer containing 60 μM DCIP, 50 mM Tris pH 8.15, 150 mM KCl, 0.1 % Triton X-100, 500 μM DHO, 100 μM CoQ0, and varied inhibitor concentrations previously incubated with the protein. At the start point, different concentrations of the inhibitors (in 100% DMSO solution) were added to the *Hs*DHODH protein solution with a fixed DMSO percent of 10%. The reaction was started with the addition of 195 μL reaction buffer without inhibitor to 5 μL of *Hs*DHODH protein solution containing inhibitor, to a final concentration of 20 nM *Hs*DHODH and varied inhibitor concentrations. As a negative control, for each incubation time, 5 μL of the enzyme solution containing 10% DMSO was added to 195 μL buffer, without the presence of the inhibitors. The reaction was monitored at 610 nm every 3 s throughout 60 s, in triplicate, for each concentration and each tested compound. The reaction buffer without protein solution was used as a blank for each point. The IC50 was determined through the graph of the relative percent of inhibition concerning the negative control versus the log of the inhibitor concentration. The dose-response curve was fit according to equation 1 using GraphPad Prism 5 software.

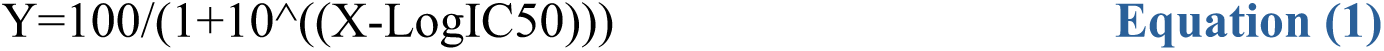

### Assessment of cytotoxicity

The cytotoxicity of compounds was assessed by MTT (3-[4,5-dimethylthiazol-2-yl]-2,5 diphenyl tetrazolium bromide) reduction assay to measure the metabolic reduction of MTT in the mitochondrial of Hep-G2 cell lines as previously described (44). Cells were cultured at 5% CO_2_ and 37°C in DMEM medium with Streptomicyn (40mg/L), supplemented with 10% heat-inactivated fetal bovine serum. Cells were seeded at a density of 10^3^ cells/well in a 96-well plate prior to incubation with a serial dilution of compounds of interest for 72h. Cells were then incubated with 3-(4,5-Dimethylthiazol-2-yl)-2,5-Diphenyltetrazolium Bromide (Sigma-Aldrich M5655) for 4h followed by formazan crystal solubilization with isopropanol and absorbance readings at OD₅₇₀. Cellular viability was expressed as a percentage relative to vehicle-treated control.

Monolayers of Calu-3 cells (2x 10^4^ cell/well) in 96-well culture plates were incubated with the compounds at different concentrations for 72 h at 37 °C at 5% CO_2_. Then, the supernatant was removed, the cells were washed with PBS and the monolayer was fixed/stained for 1 hour at 37 °C, 5% CO_²_ with methylene blue solution: HBSS + 1.25% glutaraldehyde + 0.6% methylene blue (SYNTH™). The solution was removed and the wells were washed lightly with distilled water. Subsequently, the elution solution (50% ethanol + 49% PBS + 1% acetic acid) was added and incubated for 15 min at room temperature. The volume was transferred to a flat-bottomed 96-well plate and the reading of absorbance was performed at a wavelength of 660 nm. The 50 % cytotoxic concentration (CC_50_) was calculated by performing a linear regression analysis on the dose-response curves generated from the data.

### Anti-SARS-CoV-2 assays

#### Cells, viruses, and reagents

African green monkey kidney cells (Vero, subtype E6) and human lung epithelial cell lines (Calu-3) were expanded in high glucose DMEM High Glucose with 10% fetal bovine serum (FBS; Merck), with 100 U/mL penicillin and 100 μg/mL streptomycin (Pen/Strep; Gibco) at 37°C in a humidified atmosphere with 5% CO_2_. The SARS-CoV-2 B.1 lineage (GenBank accession no. MT710714) was expanded in Vero E6 cells. Viral isolation was performed after a single passage in cell culture in 150 cm^2^ flasks with high glucose DMEM plus 2% FBS. Observations for cytopathic effects were performed daily and peaked 4 to 5 days after infection. All procedures related to virus culture were handled in biosafety level 3 (BSL3) multiuser facilities, according to WHO guidelines. Virus titers were determined as plaque-forming units (PFU/mL), and virus stocks were kept in −80°C ultralow freezers.

The SARS-CoV-2 B.1 lineage (GenBank #MT710714) was isolated in Vero E6 cells from nasopharyngeal swabs of a confirmed case. All procedures related to virus culture were handled in the BSL3 multiuser facility at Fundação Oswaldo Cruz (FIOCRUZ), Rio de Janeiro, Brazil, according to World Health Organization (WHO) guidelines (45).

#### Infections and virus titration

Calu-3 cells (2.0 × 10^5^ cells/well) in 96-well plates (Nalge Nunc Int’l, Rochester, NY, USA) were infected with a multiplicity of infection (MOI) of 0.1 for 1 h at 37 °C at 5% CO2. The inoculum was removed, and cells were incubated with treatments or not in DMEM with 10% FBS. After 48 h, the virus content in the supernatant was quantified by plaque-forming assays in Vero cells. For virus titration, monolayers of Vero E6 cells (2 x 10^4^ cell/well) in 96-well plates were infected with serial dilutions of supernatants containing SARS-CoV-2 for 1 hour at 37°C. Semi-solid high glucose DMEM medium containing 2% FBS and 2.4% carboxymethylcellulose was added and cultures were incubated for 3 days at 37 °C.

Then, the cells were fixed with 10% formalin for 2 hours at room temperature. The cell monolayer was stained with 0.04% solution of crystal violet in 20% ethanol for 1 hour. Plaque numbers were scored in at least 3 replicates per dilution by independent readers blinded to the experimental group. The virus titers were calculated by scoring for plaque-forming units (PFU/mL), and non-linear regression analysis of the dose–response curves were also performed to calculate the 50% effective concentration (EC_50_). All experiments were carried out at least three independent times, including a minimum of two technical replicates in each assay. Data were analyzed using Prism GraphPad software 9.0 (Windows GraphPad Software, San Diego, CA, USA). Values were presented as means ± standard deviations (SD).

### Water solubility determination

The solubility of synthesized compounds was determined in ultrapure water. An excess of each (1 mg) was added to beakers containing 1.5 mL of water, and dispersions were kept at room temperature under stirring (IKA, Vortex 3, Staufen, Germany) for 12 h. Then, the samples were filtered in a PVDF membrane 0.22 μm and diluted in DMSO/H2O 1:1 for absorbance reading in a UV-Vis spectrophotometer (Shimadzu, UV-1800, Kyoto, Japan). The study was conducted in triplicate for each drug.

Quartz cuvettes of 1 cm of the optical path were used for spectrophotometric quantification. Scans of known concentrations of the compounds were performed in DMSO/H2O 1:1, in the wavelength range (λ) between 220 and 600 nm, to determine the λ of maximum absorption (λmax) of each of them in the UV/Vis. The λmax was used to determine the solubility constant of the compounds from analytical calibration curves. The curves were constructed from known concentrations of the compounds in DMSO/H2O (1:1) in the concentration range from 0.1 to 10 μg/mL. Each concentration absorbance (A) was plotted on the ordered axis, and the respective concentration was on the abscissa’s axis. The least squares linear regression method was used to fit the data points. The first-order equation A = ax + b (where a is the slope, and b is the linear coefficient, given by the line intersection point with the ordinate axis) was used to convert the sample’s A in concentration (x). Linear ranges were calculated using the linear correlation coefficient (r) as the minimum acceptable criterion of r = 0.99.

### Computational session

#### Ligand Preparation

The 3D structures of the quinones were drawn using ChemDraw software v.20.1.1, imported into Maestro workspace v.12.8 (Schrödinger, LCC, New York, 2021), and prepared using the LigPrep tool (46,47). All ionization and tautomeric states were generated at pH 7.4 ± 0.5 using Epik software v.5.6 (Schrödinger, LCC, New York, 2021) (48,49). The lowest potential energy conformers and tautomers were calculated using OPLS4 (50) and retained as input for docking studies.

#### Molecular docking calculations

The *Hs*DHODH structure in complex with the inhibitor 1214 (B6U) (PDB ID: 6LP6) (51) was selected based on a similarity between the co-crystalized ligand with the quinones, and the good resolution of 1.79 Å. The 3D structure of the protein was imported into the Maestro workspace and prepared using the Protein Preparation Wizard tool (47,49) (Schrödinger, LCC, New York, 2021). Hydrogen atoms were added, according to Epik v.5.6(48,49), pKa was calculated (pH 7.4±0.5) using PROPKA (52), and energy minimization was performed using the OPLS4 force field (50). Then, a grid box of 15 Å around the Ubiquinone-binding site (51), was generated using the receptor grid generation module of the Glide v. 9.1 (Schrödinger, LCC, New York, 2021) (53,54), using the co-crystalized ligand B6U and grid coordinates *x, y,* and *z* of −31.39, 21.81, 23.20 Å, respectively. The docking was performed using Glide software v.9.1 (53,54), using the extra-precision (XP) (55), to generate 10 poses for each ligand, and the Docking XP score. Then, the LE (56) was computed according to equation 2.

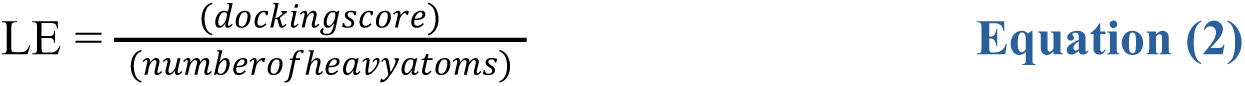

The Molecular mechanics with generalised Born and surface area solvation (MM-GBSA) (57) score was calculated using the Prime tool (58–60), according to equation 3 (57).

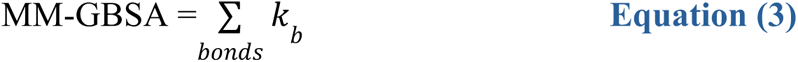

Finally, to analyze the protein–ligand interactions of the docking poses, we used PLIP server (61) and for generating the images we used Pymol software v. 2.3.0(62).

#### Molecular Electrostatic Potential

The electrostatic potential maps for the quinones were calculated using Avogadro software v. 1.2 (63,64), using extensions to generate Van der walls surfaces and color by electrostatic potential. For the *Hs*DHODH electrostatic surface, it was calculated using Pymol software v. 2.3.0 (65), with plugin APBS electrostatic, with all formal charges.

#### QSAR classification models

The QSAR classification models based on machine learning and deep learning were developed in Python v.3.9 following best practices for predictive modeling (66). To ensure the reproducibility of our computational workflow, all the source code and datasets used in this work can be found in our GitHub repository.

#### Data curation

A data set of 9,889 compounds with EC_50_ data measuring cytopathic effect induced by SARS-CoV-2 in Vero E6 cells was collected and downloaded from the ChEMBL database (ID: CHEMBL4303835) and used to develop the classification models. Briefly, all chemical structures and corresponding EC_50_ data were carefully processed and curated according to the protocols proposed by Fourches et al. (67,68) During this step, an activity threshold of 10 μM was used to distinguish into active and inactive compounds, since this value represents a good starting point for the prospective hit-to-lead optimization studies. Therefore, 107 compounds with EC_50_ ≤ 10 μM were categorized as actives, whereas 4,527 compounds with EC_50_ > 10 μM were categorized as inactives. Since the curated dataset presented a high imbalance ratio of 1:42 (107 actives vs. 4,527 inactives), it was partially balanced (ratio of 1:10, 107 actives vs. 1,070 inactives) using a linear under-sampling approach(68).

#### Machine learning model

The machine learning model was developed using Random Forest (RF) (46) algorithm implemented in Scikit-learn v.0.24.2 and Extended Connectivity Fingerprints with diameter 4 (ECFP4 with 2048 bits) implemented in RDKit v.2021.03.1. All models were built using 5-fold cross-validation approach whereas their hyperparameters were optimized using a Bayesian approach implemented in Scikit-Optimize v.0.7.4.

#### Deep learning models

In this work, FFNN (using ECFP4 fingerprints as chemical features) and message-passing neural network (MPNN)(69) architectures were explored in TensorFlow v.2.8(70) to develop deep learning models. Initially, the curated dataset was randomly divided into a training set, validation set, and test set at a ratio of 8:1:1 by Python scripts. The training set was used to build the model, the validation set for hyper-parameter optimization, and the test set was used for the model evaluation. Concerning the complexity and high computational cost of explored architectures, we used a random searching strategy(70) based on previous experience for hyperparameter settings (e.g. dropout, learning rate, batch size, activation function, L1 and L2 regularization) and message-passing(70) hyperparameters (e.g. message units, message steps, number of attention heads). The early stopping approach(70) is used to avoid overfitting and save computational resources. The performance metrics of the validation set were used for model selection.

#### Statistical validation

The predictive performance of machine learning and deep learning models was evaluated using ACC, SE, SP, positive predictive value (PPV), negative predictive value (NPV), and Matthews correlation coefficient (MCC). These metrics were calculated using the equations 4-9.

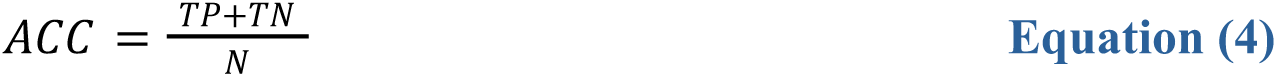

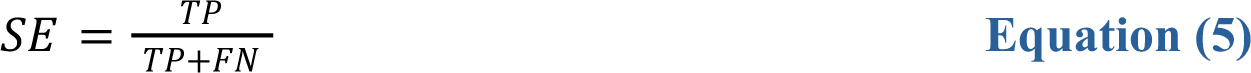

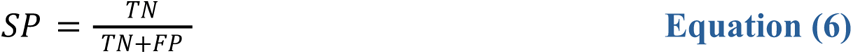

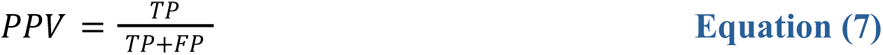

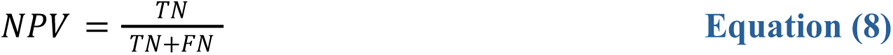

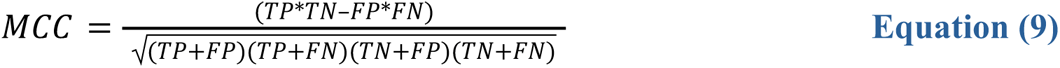

where N represents the number of compounds, FP and FN represent the number of false positives and false negatives, and TP and TN represent the number of true positives and true negatives, respectively.

#### Model Mechanistic interpretation

Predicted probability maps were generated for the best QSAR model to visualize the fragment contributions for anti-cytopathic activity against SARS-CoV-2. Here, the “weight” of a fragment (bit) was considered as the predicted-probability difference obtained when the bit is removed. The normalized weights were then used to color the atoms in a topography-like map with gray representing no changes in probability, green indicating a positive difference (i.e. probability decreases when the bit is removed) and pink indicating a negative difference(71).

#### Data and code availability

All models, datasets, and code are available on GitHub https://github.com/LabMolUFG/COVID).

## Supporting information

Supplemental data and experimental

## ACKNOWLEDGMENTS

This work has been funded by CNPq BRICS STI COVID-19 (#441038/2020-4), FAPEG (#202010267000272), São Paulo Research Foundation (FAPESP grants #2020/06190-0, #2021/10084-3, #2021/13237-5 and #2020/05369-6). CHA, TMSL, FSE, FTMC, and MCN are CNPq research fellows.

## CONFLICT OF INTEREST

The authors declare no conflict of interest.

## AUTHOR CONTRIBUTIONS

C.H.A., F.S.E. and M.C.N. coordinated, designed and supervised the project. C.H.A. and M.C.N. acquired funding for this project. B.F.G. and M.M.V. synthesized the compounds and organized the synthetic and structure elucidation experimental data. A.D.P. provided and organized the biochemical experimental data, and wrote the first draft of the manuscript. T.M.L.S. supervised the antiviral experiments. C.Q.S., J.R.T. and N.F-R. provided compounds antiviral data. S.S.M., L.V.C., M.V.F., M.F.B.S. and B.J.N. provided chemoinformatics and molecular modeling data. F.T.M. supervised the cytotoxicity experiments. J.A.L and L.C.S.A. provided and organized cytotoxicity data. R.F.V.L. supervised and B.A.M. provided compounds physico-chemical data. All authors critically reviewed and contributed to the final version of the paper.

## ABBREVIATIONS

ACC: accuracy
BSL3: biosafety level 3
DHO: dihydroorotate
DHODH: dihydroorotate dehydrogenase
DNA: deoxyribonucleic acid
FBS: fetal bovine serum
FDA: Food and drug administration
FFNN: feedforward neural network
FMN: flavin mononucleotide
HsDHODH: human dihydroorotate dehydrogenase
LE: ligand Efficiency
MCC: Matthews correlation coefficient
MM-GBSA: Molecular mechanics with generalized Born and surface area solvation
MPNN: message-passing neural network
MTT: (3-[4,5-dimethylthiazol-2-yl]-2,5 diphenyl tetrazolium bromide)
NPV: negative predictive value
NQ: 2-hydroxynaphthoquinone
ORO: orotate
PFU: plaque-forming units
PPV: positive predictive value
QSAR: quantitative structure-activity relationship
RF: random Forest
RNA: ribonucleic acid
SE: sensitivity
SmDHODH: Schistosoma mansoni
DHODH SP: specificity
UMP: uridine monophosphate

## Notes

### Competing Interest Statement

The authors have declared no competing interest.

