## Supplemental data and experimental for "Unveiling the antiviral capabilities of targeting Human Dihydroorotate Dehydrogenase against SARS-CoV-2"

### SUPPLEMENTARY FIGURES AND TABLES

**FIGURE S1.** Superposition of all quinones docked in the *Hs*DHODH structure (gray) with the crystal structure of DHODH co-crystalized with brequinar (blue) (PDB ID: 1D3G), and teriflunomide (PDB ID: 1D3H), in pink.

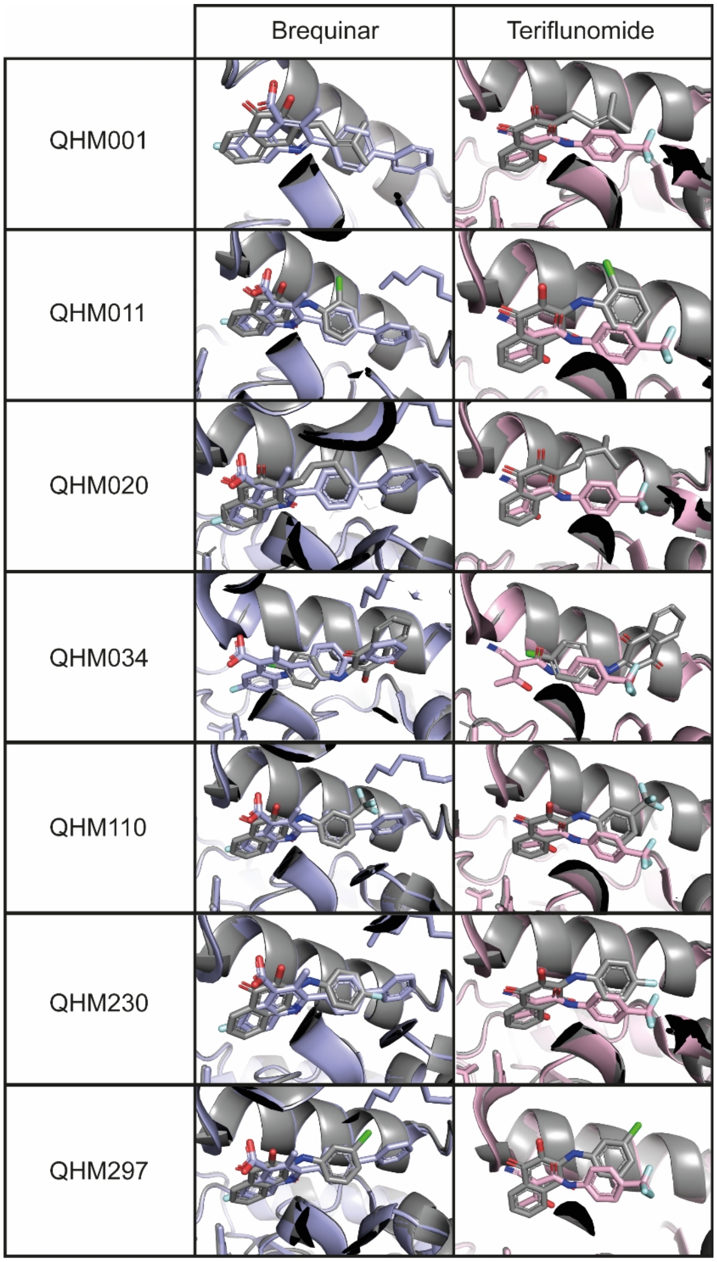

**FIGURE S2.** 3D Conformations obtained by docking for HsDHODH in complex with QHM001 (A) and QHM110 (B). *Hs*DHODH is represented by lines with the molecular electrostatic surface, and the ligand as stick. Protein and ligand colored by elements: oxygen red, fluorine sky blue, nitrogen blue and carbon yellow for QHM001, green for QHM110 and pink for the protein.

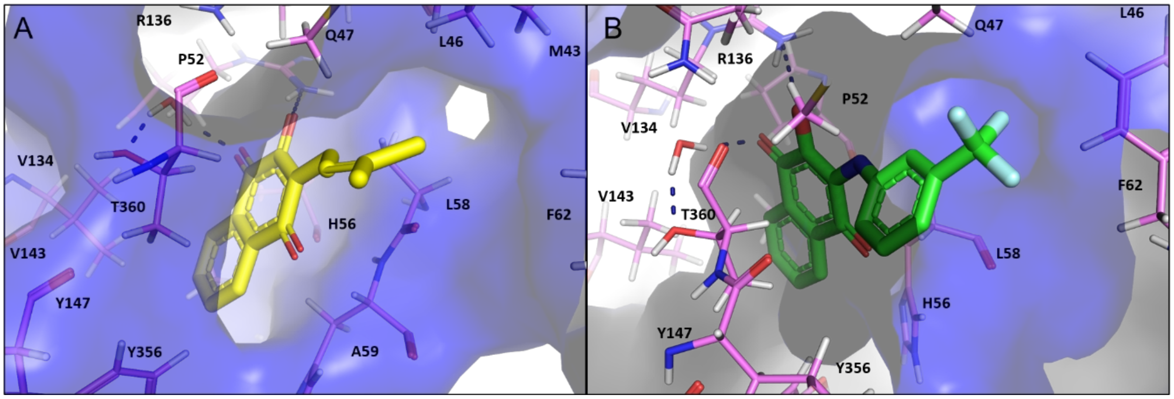


**FIGURE S3.** Anti SARS-CoV-2 dose-response curves for all the tested compounds, including quinone-based compounds and the experimental controls**.**

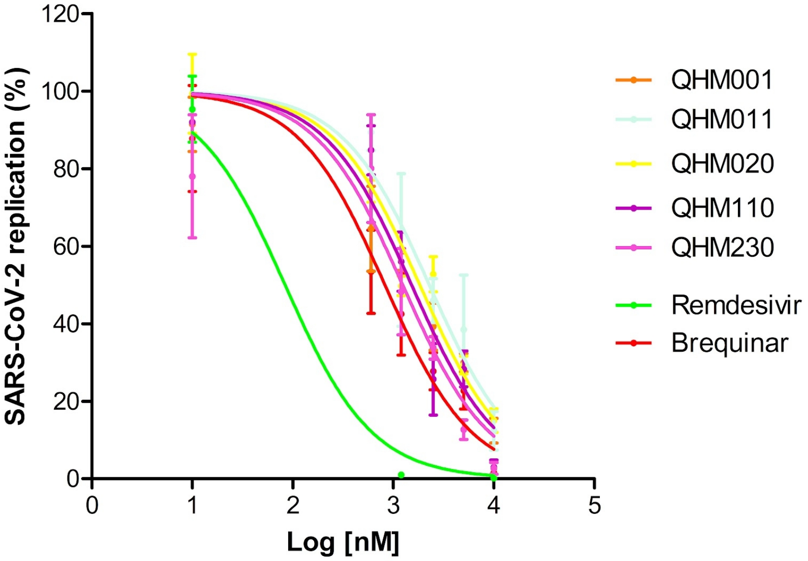

**TABLE S1.** Molecular docking results of the best ligands with experimental data ($IC_{50}$), with Glide XP Score, Ligand Efficiency (LE), and MM-GBSA score.

| Ligand | Glide XP Score (kcal/mol) | LE (kcal/mol) | MM-GBSA (kcal/mol) |
|---|---|---|---|
| QHM110 | -10.99 | -0.46 | -66.67 |
| QHM011 | -9.84 | -0.47 | -67.18 |
| QHM230 | -9.57 | -0.45 | -66.08 |
| QHM297 | -9.26 | -0.44 | -65.94 |
| QHM001 | -9.00 | -0.50 | -44.14 |
| QHM020 | -7.28 | -0.40 | -41.33 |
| QHM034 | -9.34 | -0.45 | -56.64 |

**TABLE S2.** Experimental water solubility for the quinone compounds.

| Identification of compounds | Solubility (µg/mL) | Solubility (µM) |
|---|---|---|
| QHM0001 | 4.5 ± 0.4 | 18 ± 2 |
| QHM0011 | 0.8 ± 0.2 | 2.7 ± 0.6 |
| QHM0020 | 2.7 ± 0.5 | 11 ± 2 |
| QHM0034 | 31 ± 3 | 102 ± 9 |
| QHM0110 | 20 ± 1 | 58 ± 3 |
| QHM0230 | 1.5 ± 0.1 | 5.2 ± 0.5 |
| QHM0297 | 13 ± 2 | 43 ± 6 |

**TABLE S3.** Statistical characteristics of best QSAR models for the training, validation and test sets.

| MODEL | DATA SET | *Metrics* | | | | | |
|---|---|---|---|---|---|---|---|
| | | ACC | SE | SP | PPV | NPV | MCC |
| FFNN | Train | 0.94 ± 0.02 | 0.69 ± 0.21 | 0.98 ± 0.01 | 0.72 ± 0.08 | 0.96 ± 0.02 | 0.62 ± 0.19 |
| | Test | 0.94 ± 0.04 | 0.74 ± 0.27 | 0.96 ± 0.03 | 0.62 ± 0.23 | 0.97 ± 0.0 | 0.64 ± 0.25 |
| MPNN | Train | 0.93 ± 0.01 | 0.74 ± 0.09 | 0.94 ± 0.01 | 0.31 ± 0.08 | 0.99 ± 0.01 | 0.44 ± 0.04 |
| | Test | 0.90 ± 0.03 | 0.40 ± 0.21 | 0.92 ± 0.04 | 0.19 ± 0.09 | 0.97 ± 0.01 | 0.22 ± 0.03 |
| RF | Train | 0.92 ± 0.02 | 0.53 ± 0.16 | 0.95 ± 0.04 | 0.59 ± 0.20 | 0.95 ± 0.01 | 0.50 ± 0.10 |
| | Test | 0.92 ± 0.02 | 0.53 ± 0.16 | 0.95 ± 0.04 | 0.59 ± 0.20 | 0.95 ± 0.01 | 0.25 ± 0.14 |

### CHEMISTRY SESSION

Considering the strategy of ring bioisosterism and molecular simplification based on the atovaquone structure, we synthesized epoxide **2** to carry out a ring-opening reaction with several anilines toward substituted 2-hydroxynaphthoquinones (adapted from Calil et al). The final compounds were synthesized using isopropyl alcohol as solvent, which were obtained in low to good yields (10-60%).

Among lapachol derivatives (series B), the reduced lapachol was afforded using palladium catalyzed reductive hydrogenation under 100psi pressure.

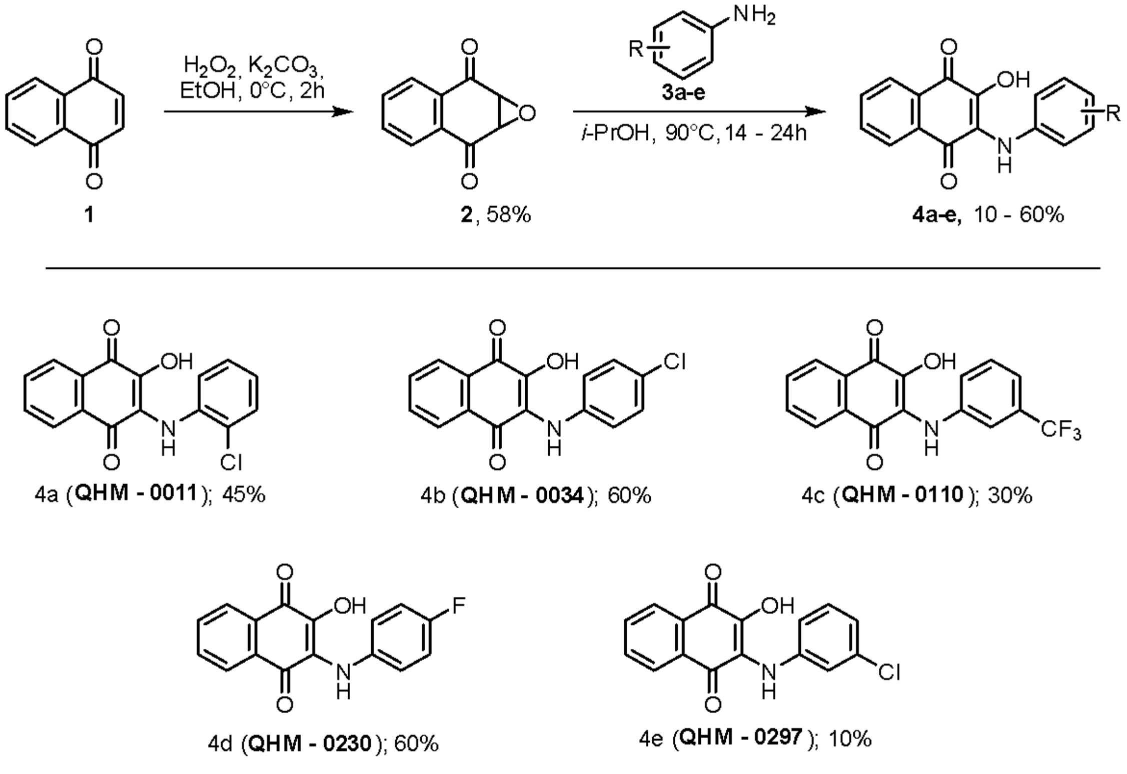


**Scheme S1. Synthesized quinone-based compounds 2-hydroxy-3-amino-1,4-naphthoquinones (series C and D).**

### Material and Methods

**General remarks for chemistry:** All commercially available reagents and solvents were used in the reactions without further purification. Yields refer to isolated and purified products, unless otherwise noted.

Reactions were monitored by thin layer chromatography (TLC) and visualized under UV (Ultraviolet) light at 254 and 365 nm. Column chromatography was performed using silica gel 60 (70-230 mesh) or using Flash chromatography purification system CombiFlash® Rf$^{+}$. $^{1}$H and $^{13}$C NMR spectra were recorded at 300 MHz and 75 MHz, respectively. Chemical shifts were referenced to the deuterated solvent (i.e., for CDCl3, δ = 7.26 and 77.16; for DMSO-d6, δ = 2.50 and 39.52, for $^{1}$H and $^{13}$C NMR, respectively) and are reported in parts per million (ppm, δ). Coupling constants (J) are stated in Hz using the splitting abbreviations: s, singlet; d, doublet; t, triplet; m, multiple. High-resolution mass spectra (HRMS) were measured by a TOF spectrometer, using electrospray ionization (ESI). The purity of synthesized compounds was determined by LCMS (Liquid Chromatography – Mass Spectra), using a Kinetex C18 column (150 x 4,6mm), and 5µM as the size of the particle at 254nm or by $^{1}$H qNMR, and the samples showed more than 95% of purity.

***Synthesis of 1,4-naphtoquinone-based compounds*. General procedure for the synthesis of 1,4-naphtoquinone epoxide 2 (1a,7a-dihydronaphtho[2,3-b]oxirene-2,7-dione).**

To a solution of 1,4-naphthoquinone (5.0 g, 31.6 mmol, 1 equiv) in ethanol (150 mL) at 0°C, hydrogen peroxide (20% in water – 21,50 mL, 126,04 mmol, 4 equiv) was added dropwise, followed by slow addition of potassium carbonate (8,74 g, 63,23 mmol, 2 equiv) in distilled water. The reaction mixture was stirred at 0 °C, covered from light, for 2h and quenched afterwards by addition of water (150ml). The precipitated compound mixture was filtered and partitioned with ethyl acetate (3x50 mL). The organic layer was collected, dried over $Na_2SO_4$, and concentrated under reduced pressure. The crude material was quickly purified by column chromatography on silica gel with hexanes/ethyl acetate (9:1) to afford the desired product as white crystals, which was readily stored at -20°C and protected from light (3,20g, 58,12%). $^1$H NMR (300 MHz, DMSO-d6) δ 7.95-7.85 (m, 4H), 4.17 (s, 2H); $^{13}$C NMR (75 MHz, DMSO-d6) δ 190.7, 134.7, 131.4, 126.5, 55.3. The spectral data are in accordance with those reported in the literature (1). Purity: >99.90% (assessed by LC/MS).

**General procedure for the synthesis of 1,4-naphtoquinone-based compounds with anilines.**

To a suspension of **2** (1 equiv) in isopropyl alcohol (10ml), the desired aniline (1,2 equiv) was added, and the mixture was stirred at 90°C for 14-24h (until the indication of no further amount of **2** or the increasing amount of lawsone as a side product). After this time, the reaction mixture was cooled down and partitioned with ethyl acetate. The organic layer was collected, dried over $Na_2SO_4$ and concentrated. The products were purified over automatic silica gel column (Combiflash rf+) with hexane/dichloromethane gradient to afford the desired products (Yields shown on Scheme 1).

**General procedure for the synthesis of 2-hydroxy-3-isopentylnaphthalene-1,4-dione (Reduced lapachol).**

To a solution of lapachol (242 mg, 1 mmol, 1 equiv) in ethyl acetate was added Pd/C (10 mg) and the mixture was stirred under $H_2$ atmosphere (100psi) at room temperature for 2 h.

The mixture reaction was concentrated to dryness and the residue was purified by column chromatography on silica gel with hexane/dichloromethane (95:5) to afford the desired product as yellow solid (195 mg, 83%). $^{1}$H NMR (300 MHz, $CDCl_3$) d 8.11 (dd, *J* = 7.7, 1.1 Hz, 1H), 8.07 (dd, *J* = 7.5, 1.3 Hz, 1H), 7.75 (td, *J* = 7.5, 1.4 Hz, 1H), 7.67 (td, *J* = 7.5, 1.3 Hz, 1H), 7.29 (s, 1H), 2.64 - 2.56 (m, 2H), 1.69 - 1.54 (m, 1H), 1.47 - 1.34 (m, 2H), 0.97 (s, 3H), 0.94 (s, 3H); $^{13}$C NMR (75 MHz, $CDCl_3$) 184.8, 181.6, 153.0, 134.9, 133.1, 133.0, 129.5, 126.8, 126.2, 125.2, 37.3, 28.4, 22.5, 21.5. Data in accordance with the literature (1). Purity: 99.97% (assessed by LC/MS).

*2-((2-chlorophenyl)amino)-3-hydroxynaphthalene-1,4-dione* (**4a – QHM0011**)

151 mg (45%); dark purple solid. $^{1}$H NMR (300 MHz, DMSO-$d_6$) δ 10.66 (s, 1H), 8.00-7.95 (m, 2H), 7.82-7.75 (m, 2H), 7.40 (dd, *J* = 6.9, 1.2 Hz, 1H), 7.20 (td, *J* = 7.7, 1.7 Hz, 1H), 7.10 (s, 1H), 6.95 (td, *J* = 7.5, 1.4 Hz, 1H), 6.86 (dd, *J* = 8.1, 1.6 Hz, 1H); $^{13}$C NMR (75 MHz, DMSO-$d_6$) d 219.2, 217.2, 180.1, 175.4, 171.5, 171.4, 168.3, 168.1, 166.4, 164.5, 163.4, 163.2, 162.6, 161.1, 160.0, 159.4. Data in accordance with the literature (1). Purity: 98.64% (assessed by LC/MS).

*2-((4-chlorophenyl)amino)-3-hydroxynaphthalene-1,4-dione* (**4b – QHM-0034**):

180 mg (60%); dark purple solid. $^{1}$H NMR (300 MHz, DMSO-$d_6$) δ 10.47 (s, 1H) 8.16 (s, 1H), 7.95 (m, 2H), 7.77 (m, 2H), 7.20 (d, *J* = 8.5 Hz, 2H), 6.85 (d, *J* = 8.6 Hz, 2H); $^{13}$C NMR (75 MHz, DMSO-$d_6$) d 181.9, 179.5, 142.4, 140.5, 133.7, 133.6, 130.9, 130.5, 127.5 (2), 125.7, 125.4, 125.2, 123.8, 120.7 (2). Data in accordance with the literature (1). Purity: >95% (assessed by $^{1}$H qNMR).

*2-hydroxy-3-((3-(trifluoromethyl)phenyl)amino)naphthalene-1,4-dione* (**4c – QHM-0110**)

110 mg (30%); dark purple solid. $^1$H NMR (300 MHz, DMSO-$d_6$) δ 10.70 (s, 1H), 8.33 (s, 1H), 8.02 - 7.93 (m, 2H), 7.80-7.77 (m, 2H), 7.37 (t, *J* = 7.9 Hz, 1H), 7.16 (s, 1H), 7.15-7.04 (m, 2H); $^{13}$C NMR (75 MHz, DMSO-$d_6$) d 182.3, 180.1, 143.7, 142.8, 134.2, 134.1, 131.3, 130.9, 129.1 (d, *J* = 31.3 Hz), 129.1, 126.1, 125.9, 125.0, 124.8 (d, *J* = 272.3 Hz), 122.7, 116.3 (d, *J* = 4.2 Hz), 115.3 (d, *J* = 3.9 Hz). Data in accordance with the literature (1). Purity: 95.29% (assessed by LC/MS).

*2-((4-fluorophenyl)amino)-3-hydroxynaphthalene-1,4-dione* (**4d – QHM-0230**);

169 mg (60%); dark purple solid. $^1$H NMR (300 MHz, DMSO-$d_6$) δ 10.21 (s, 1H), 8.03 (s, 1H), 7.97-7.93 (m, 2H), 7.80-7.70 (m, 2H), 7.02 (dt, *J* = 12.0, 2.9 Hz, 2H), 6.89 (m, 2H). $^{13}$C NMR (75 MHz, DMSO- $d_6$) δ 182.0, 179.1, 157.0 (d, *J* = 236.9 Hz), 140.8, 137.5, 133.8, 133.4, 130.7, 130.6, 126.0, 125.6, 125.3, 121.2 (d, *J* = 7.9 Hz), 114.2 (d, *J* = 22.3 Hz). Data in accordance with the literature (1). Purity: 97.81% (assessed by LC/MS).

*2-((3-chlorophenyl)amino)-3-hydroxynaphthalene-1,4-dione* (**4e – QHM-0297**):

30 mg (10%); dark purple solid. $^1$H NMR (300 MHz,DMSO-$d_6$) δ 10.61 (s, 1H), 8.17 (s, 1H), 7.93 (s, 2H), 7.82-7.34 (m, 2H), 7.17 (t, *J* = 7.9 Hz, 1H), 6.88-6.77 (m, 3H); $^{13}$C NMR (75 MHz, DMSO-$d_6$) δ 181.9, 179.7, 143.2, 143.1, 133.7 (2C), 132.4, 130.9, 130.5, 129.2, 125.7, 125.4, 124.6, 119.4, 118.2, 117.3. Data in accordance with the literature (1). Purity: 95,15% (assessed by $^1$H qNMR).
